# Using coexpression to explore cell-type diversity with the fcoex package

**DOI:** 10.1101/2021.12.07.471603

**Authors:** Tiago Lubiana, Helder Nakaya

## Abstract

Here, we present the fcoex package, which infers coexpression from scRNA-seq data and yields multiple, overlapping classes of cells based on coexpression modules. The tool extends the current scRNA-seq toolbox, providing a multi-hierarchy view on cell functionality and enabling the development of more complete cell atlases. Single-cell RNA sequencing (scRNA-seq) captures details of the cellular landscape, basing a fine-grained view on biological processes. Current pipelines, however, are restricted to single-label perspectives, missing details of the classification landscape. In the *pbmc3k* blood cell dataset, fcoex detects known classes, like antigen-presenting cells and a new theoretical group of cells, marked by the expression of FCGR3A (CD16). Fcoex extends the current scRNA-seq toolbox, providing a multi-hierarchy view on cell functions as a tool to develop complete cell type atlases.

**Availability and Implementation:** Fcoex is written in R and openly available in Bioconductor (https://bioconductor.org/packages/fcoex/).

**Supplementary information:** Supplementary data is available at the end of the manuscript. Source code for analysis is available at https://github.com/csbl-inovausp/fcoex_analysis;

## Introduction

Since the 17th century, science has increasingly appreciated how cells act as building blocks of life. (Mazzarello, 1999) The advent of molecular profiling has offered new opportunities to tap into the diversity of cell behaviours in the past decades. Among the molecular methods, single-cell RNA sequencing (scRNA-seq), in particular, has been rapidly growing as a method for deep characterisation of cell types and states, and its use is central for the Human Cell Atlas Project. (Regev et al., 2017)

Current scRNA-seq data analyses often rely on unsupervised clustering of cells. For that, bioinformaticians tailor parameter sets to a target resolution, i.e., the level of detail used to detect cell identities. (Luecken and Theis, 2019; Regev et al., 2017) When the clustering is finished, the groups of cells are annotated with class labels, representing the underlying biology in a language we can understand. (Clarke et al., 2021)

Single clusters, however, are not limited to a single biological function. Even single cells display simultaneously genetic programs that dictate both identity and activity. (Kotliar et al., 2019) For example, B cells act as lymphocytes alongside NK and T cells (Meehan et al., 2013) but can present MHC II antigens like dendritic cells and macrophages. (Adler et al., 2017) That multifunctionality is not unique to B cells; different cell types often share effector genes (Arendt et al., 2016). Standard scRNA-seq clustering can only display a fraction of that diversity.

Here, we focus on the challenge of identifying the multiple functional classes of cells present in scRNA-seq data. The fcoex framework uses gene co-expression to infer parallel cell clusters, emphasising the multifunctional nature of cells. By applying it to a well-known dataset of blood cells, pbmc3k, we demonstrate that fcoex recovers biologically relevant superclasses and seamlessly adds value to standard scRNA-seq data analyses.

### The fcoex method

The fcoex method is designed for application right after a standard scRNA-seq clustering step (Fig. 1A). The cluster assignments convey information about the relations between cells to the algorithm and help to guide feature selection. Then, the package selects global marker genes specific to 1, 2, or more previously defined clusters. It ranks markers by symmetrical uncertainty, a non-linear correlation metric based on classical Shannon entropy. (Yu and Liu, 2003)

**Fig. 1.**
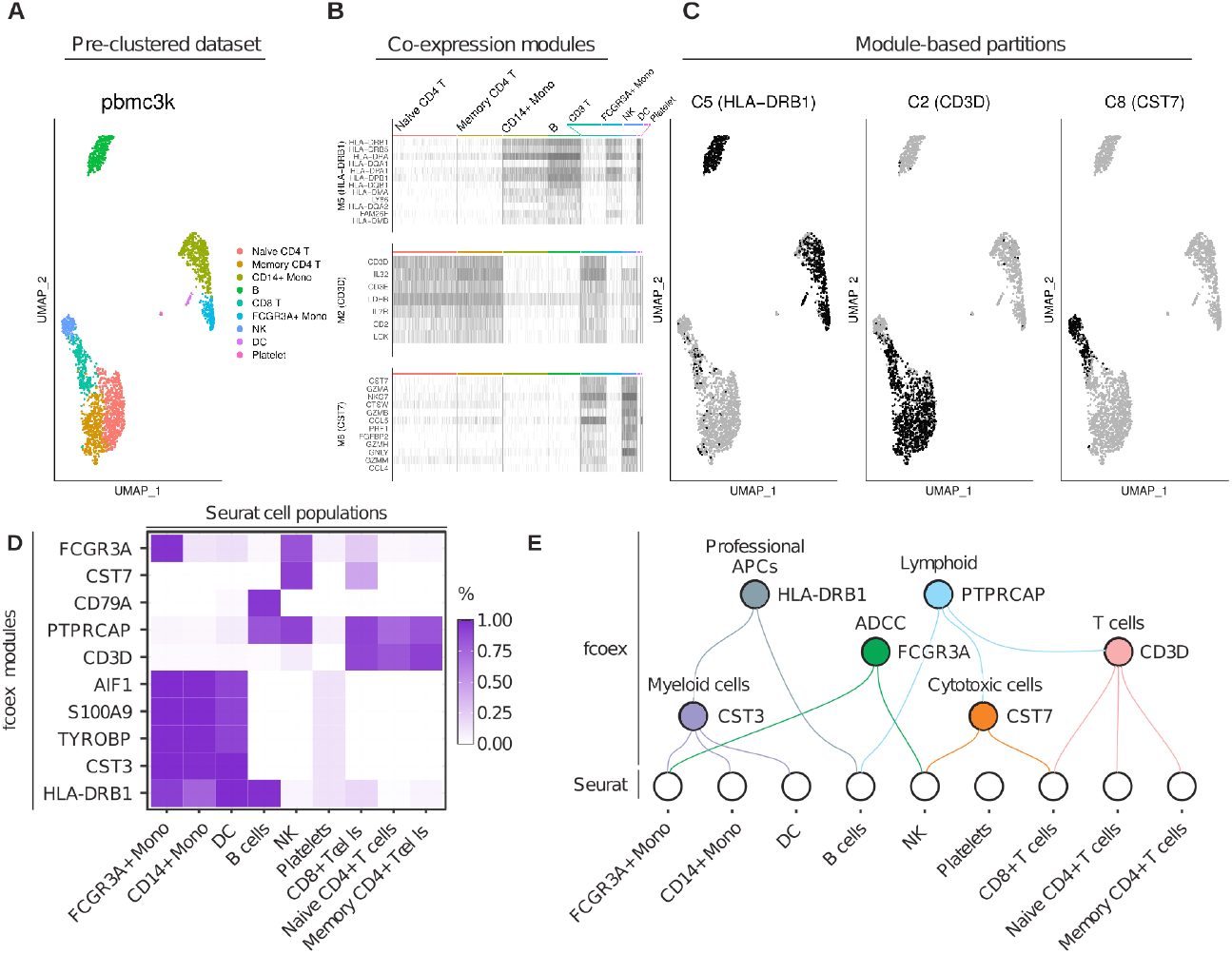
Reclustering based on co-expression modules sheds light on cell diversity. The pre-labelled pbmc3k dataset, with original, SeuratData labels. B) fcoex coexpression modules capture patterns across PBMC populations. C-D) Module-based reclustering shows groupings of original clusters, visible both via UMAP projection and cluster x cluster heatmap. E) Upper-level, multilayer classification of the pbmc3k dataset based on fcoex modules. APCs: Antigen Processing Cells; ADCC: Antibody-dependent cellular cytotoxicity

To find co-expression modules, fcoex inverts the FCBF feature selection algorithm, and instead of removing redundancy, it selects redundant (co-expressed) gene expression patterns. (see Supplementary Data for details). The default gene coexpression modules yielded by the pipeline are small by design (10s of genes per module) to facilitate manual exploration of the coexpression landscape. Each module has one “header” gene, which expression pattern better represents all the genes in the module.

Fcoex treats each module as a gene set to find cell populations, using only their expression to re-cluster cells. The new classifications are based, thus, on the genes (and functions) captured by each co-expression module. The multiple module-based clusters serve as a platform for exploring the diversity of the dataset and identifying upper cell classes, grouping cells by common functions.

### Multi-hierarchies of blood types

To validate the fcoex pipeline, we selected the well-known pbmc3k dataset from the SeuratData R package, which contains around 2700 peripheral blood mononuclear cells (PBMC) with previously-defined cluster labels.

The standard fcoex pipeline detected nine modules that capture different parts of the cellular diversity in the dataset. For example, module M8 contains cytotoxicity genes, as PRF1 and GZMA, and splits the dataset into cytotoxic (NK and CD8) and non-cytotoxic cells. M2 (CD3D) splits the dataset into T-cells and non-T-cells. M5 (HLA-DRB1) groups together monocytes, B cells, and dendritic cells, all known antigen-presenting cells (APC). (Fig. 1B-E) The classifications provided by fcoex are easily reintegrated to Seurat to power visualisations and get differentially expressed markers, providing more genes for the analysis, if desired.

In general, fcoex clusters combined biologically similar cell types of the original dataset. The clusterings help to explore and classify cells by function (Fig. 1E). Even in a well-studied dataset, fcoex provided a new light on the shared functionality of some NK cells and macrophages: they both markedly express the CD16-coding gene FCGR3A, whose product is a key player in antibody-dependent cellular cytotoxicity (ADCC). (Yeap et al., 2016) Thus, a complete functional classification of blood cells might want to include a “professional ADCC cells” class.

## Conclusion

Here we presented fcoex, a R/Bioconductor package for co-expression-based reclustering of single-cell RNA-seq data. We note that other methods are increasingly available for co-expression analysis of single cells. The monocle R package (Cao et al., 2019), widely used for pseudotime analysis, has implemented algorithms for detecting co-expression modules and WGCNA, widely used in bulk transcriptomics, has also been applied to scRNA-seq. (Langfelder and Horvath, 2008; Cardozo et al., 2019) In principle, any of those algorithms could be used as input for our framework. Of note, fcoex modules are generally smaller and simpler to explore, making it a sensible first-pass approach to explore the multi-layered diversity in single-cell transcriptomics datasets.

Regardless of the algorithm used to identify coexpression, the main goal of the fcoex pipeline is to use the modules to find biologically relevant populations. That framework allows a multi-hierarchy view of cell types, in which each cell type is related to multiple, overlapping cell classes, as seen in the Cell Ontology (Diehl et al., 2016; Osumi-Sutherland et al.,, 2021). By doing so, fcoex offers new avenues to explore data-driven classifications of cells, aligning itself with the challenges of the Human Cell Atlas and building ontologies of cell types in the single-cell era.

## Acknowledgements

We would like to thank Diógenes Lima, Pedro Russo, Gustavo Ferreira and Lucas Cardozo for their contributions to software development and Youvika Singh for helping to discuss and validate the results. We also thank all current and former members of the Computational Systems Biology Laboratory for discussions and feedback.

## Author Contribution Statement

TL and HN conceived the idea. TL wrote the software. HN supervised the project. TL and HN wrote the manuscript.

## Funding

This work was supported by the São Paulo Research Foundation (FAPESP) grants #2018/10257-2 and #2019/26284-1

## Publication Statement

This manuscript was formatted for publication in the *Application Notes* format at *Bioinformatics* and desk-rejected. The manuscript will *not* be sent to publication in other peer-reviewed journals. Nevertheless, the authors welcome reviews at any platform of open peer review (i.e. https://prereview.org/) and share a commitment to provide updated versions of the manuscript in light of any critical suggestions.

## Conflict of Interest

none declared.

## Supplementary Methods

### Preprocessing

#### pbmc3k dataset preprocessing

The *pbmc3k* dataset was loaded into R 4.1.0 as a Seurat object from the SeuratData package. The expression matrix in the “data” slot and the labels in the “Idents” slot as input for creating the *fcoex* object.

### Gene expression discretization

As the original Fast Correlation-Based Filter (Yu and Liu, 2003) algorithm was constructed to deal with discrete data, we had to discretize gene counts. We chose as a discretization metric a min-max-percent approach. For each gene, we took the lowest and the highest normalized value across the cells. and set a threshold at 25% of the max-min range. All the values below this threshold were considered “OFF,” and all above were considered “ON”.

### Identification of *fcoex* modules

#### Filtering genes by correlation to labels

After the discretization step, genes were ranked by their correlation to the labels previously assigned by the Seurat experts. We used the symmetrical uncertainty (SU), which is a nonlinear correlation metric, to rank genes. SU is a variation of mutual information that maps the values between 0 (worst) and 1 (best), and accounts for differences in entropy ranges that arise when variables have a different number of classes (number of labels and number of gene classes). All downstream steps were performed only with the genes that showed an SU over 0.1 with the labels (*fcoex*’ default).

#### Building the coexpression network

Using the genes selected by SU, fcoex first builds a full, weighted graph, where nodes represented genes and edges represent the identified SU correlation between the expression patterns of the genes. This adjacency matrix is then trimmed, and edges between nodes Yi and Yj are removed from the network iff SU(Yi, Yj) < SU(Yi, L) or SU(Yi, Yj) < SU(Yj, L). Y represents the set of expression vectors for each gene, and L represents the vector containing the cluster assignments from Seurat.

Coexpression modules are inferred from the trimmed matrix using a seed-based heuristic. Seeds for the modules are obtained via the FCBF algorithm (Yu and Liu, 2003) and consist of genes predominantly correlated to the cell classes. Each module M is composed of one module seed (x) and all the genes associated to it in the trimmed network. The modules formed in that way are fuzzy, and each gene might belong to several modules.

### Reclustering of cells

We used the *recluster* function of *fcoex* to reclassify the original dataset based on each coexpression function. The *recluster* function uses the gene sets in each co-expression community to subset the original expression table. This reduced table contains only the expression values of the in the module, and distances between cells are calculated in this reduced space. We used *fcoex* default: we calculated dissimilarities using the Manhattan distance and hierarchically clustered cells using the *ward*.*D2* metric as implemented in the *hclust* function of the R package *stats*. Two groups of cells were retrieved from each clustering (the k parameter was set to 2).

### Visualization

For visualizing the results, we used the UMAP dimensional reduction and the *DoHeatmap* and *DimPlot* functions provided in the *Seurat* package v4.0.2 (Hao *et al*., 2021) and the R package *ggplot2*.

## Code availability

The *fcoex* package, which performs the coexpression analysis, is available at http://bioconductor.org/packages/fcoex/. The discretization and feature selection algorithms are available in a second package, FCBF (http://bioconductor.org/packages/FCBF/). All the analyses performed for this work are available at https://github.com/csbl-inovausp/fcoex_analysis.

## Supplementary Figures

**Supplementary Figure 1:**
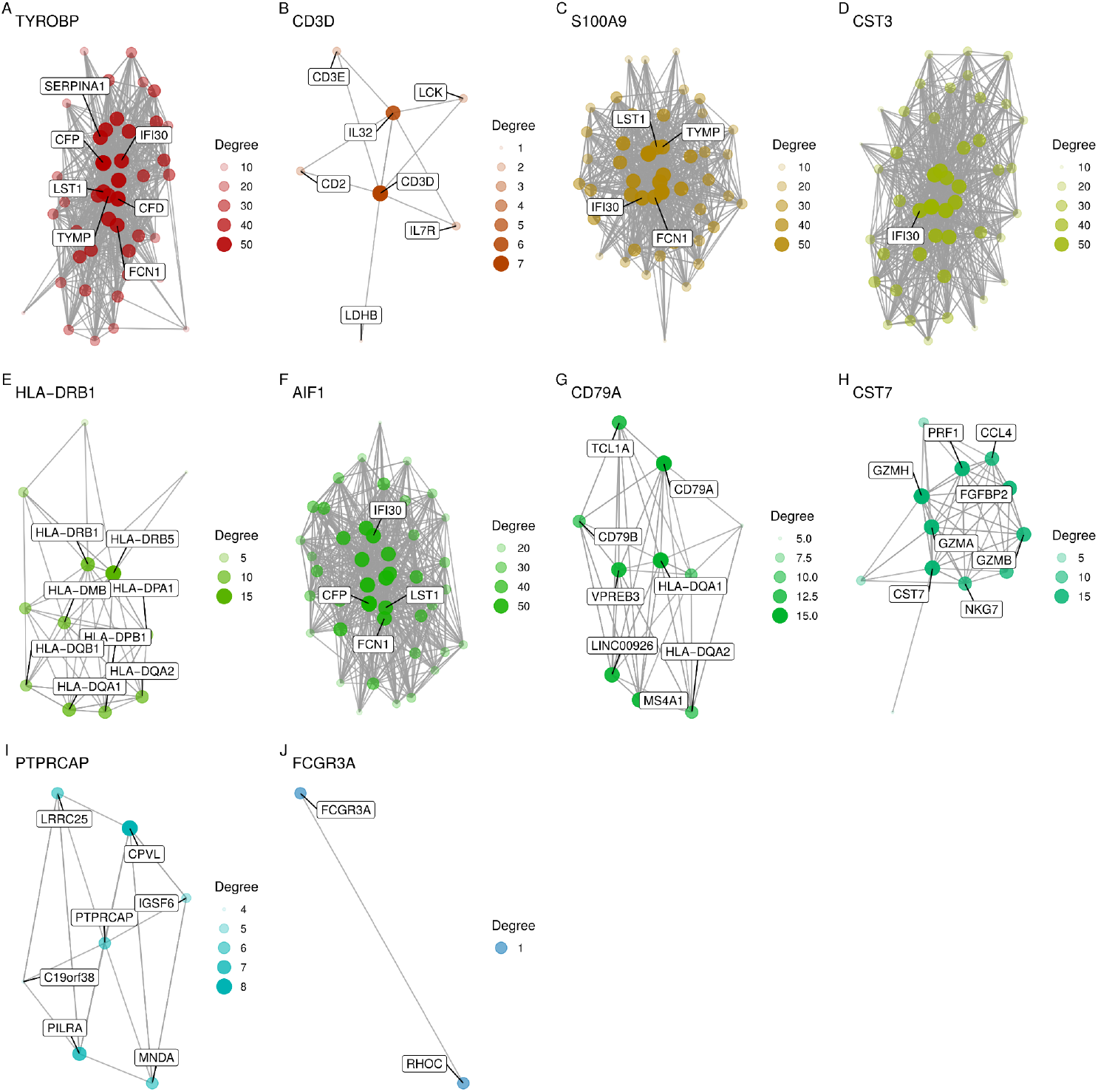
Fcoex generates small, interpretable modules. A-J) The 10 modules generated by the default *fcoex* pipeline for the *pbmc3k* dataset. The plots were generated via the *get_nets* function of the *fcoex* package. The code for the function was based on the CEMiTool package. (Russo *et al*., 2018)

**Supplementary Figure 2:**
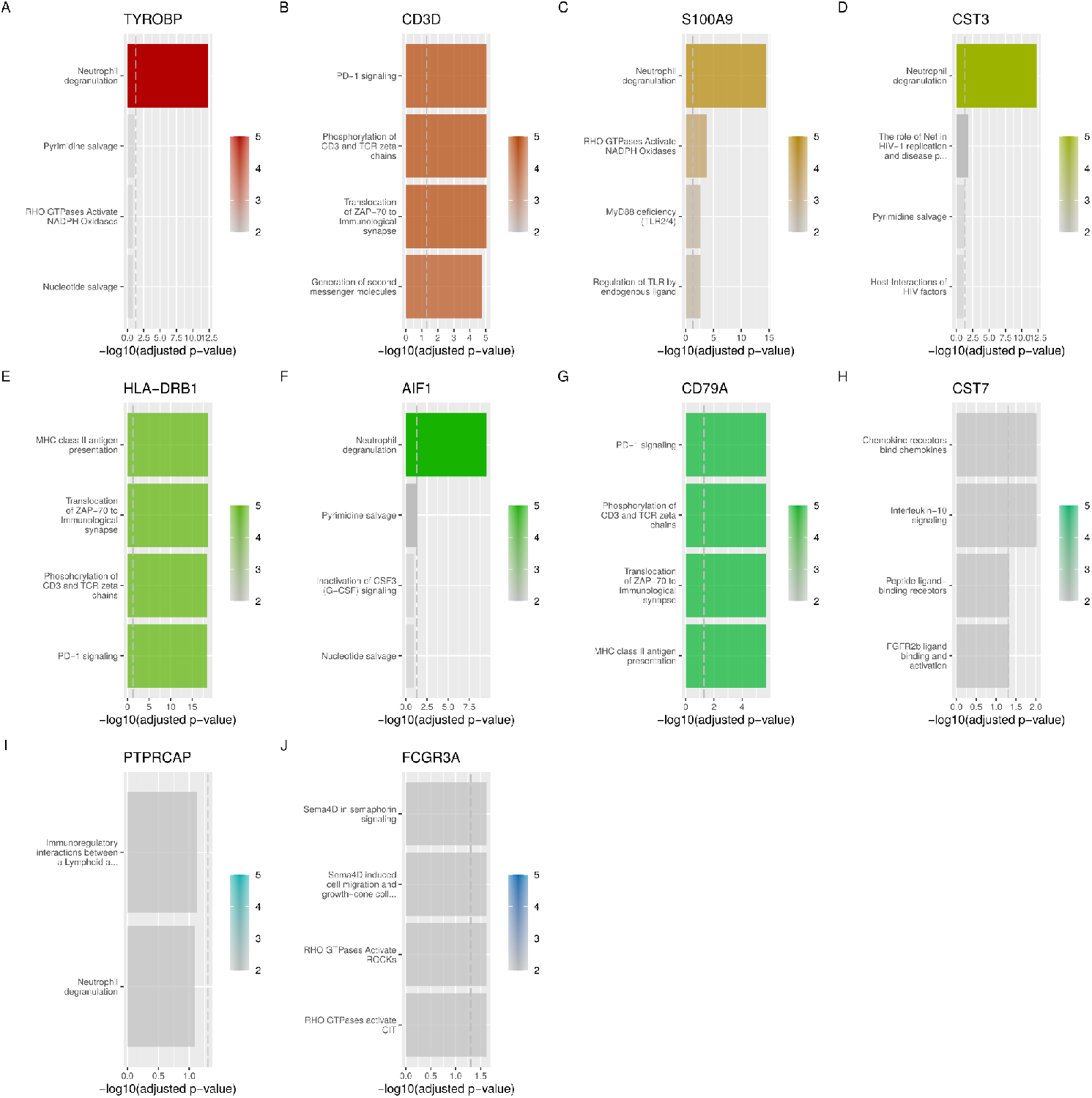
Fcoex modules are enriched for relevant biological pathways. A-J) The up-to-4 more enriched Reactome pathways (Jassal *et al*., 2020) for each of the 10 modules generated by the default *fcoex* pipeline for the *pbmc3k* dataset. Even though the number of genes per module is small, in most cases it is possible to see enrichment for biological functions associated with subsets of PBMCs. The enrichment calculation was done via the functions *mod_ora* and *plot_ora* of the *fcoex* package. The code for both functions was based on the implementation of the CEMiTool package. (Russo *et al*., 2018)

**Supplementary Figure 3:**
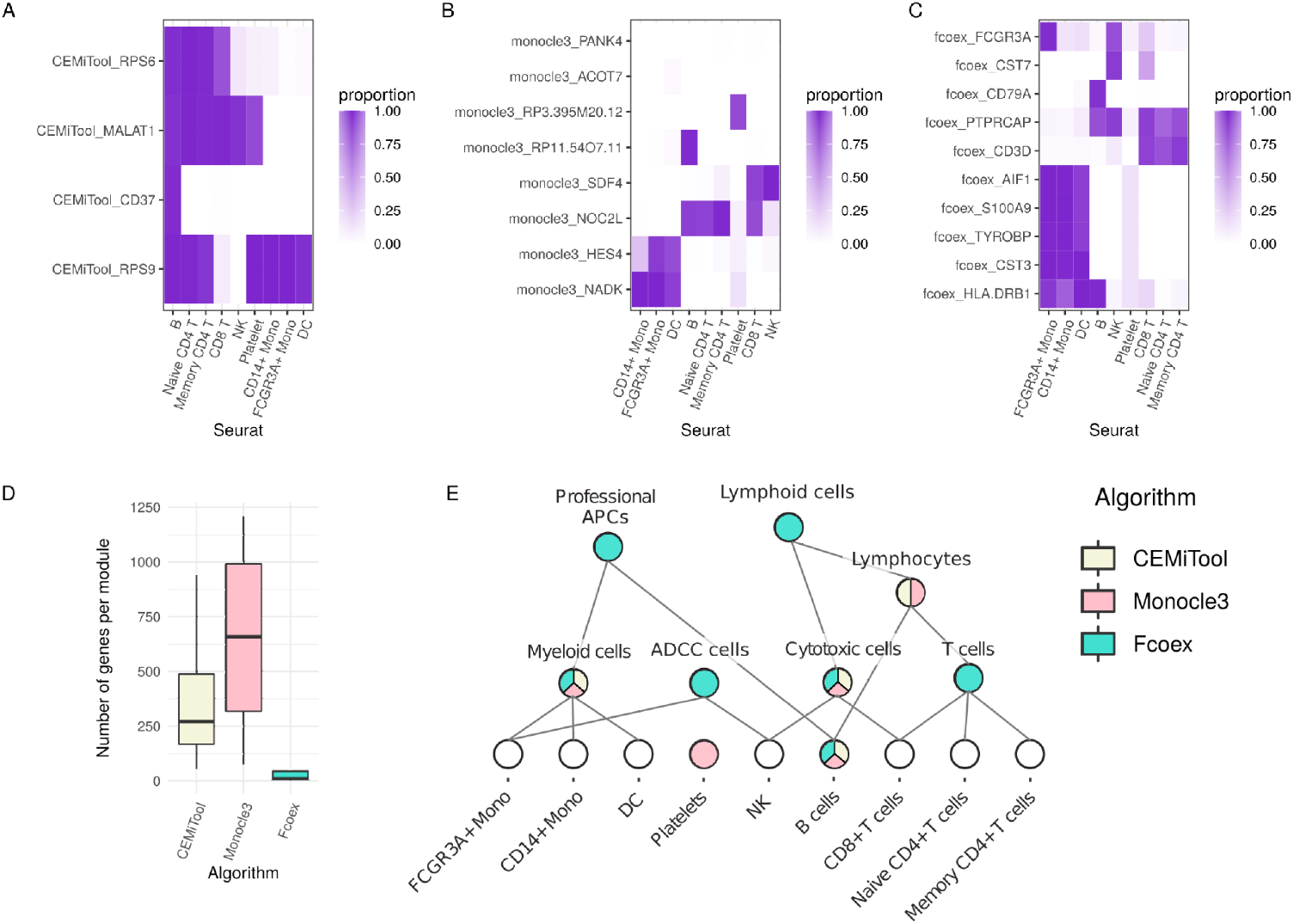
Different co-expression algorithms highlight different facets of cell diversity. A-C) Reclustering of the *pbmc3k* dataset based on modules obtained via the default pipelines for inference of coexpression modules: CEMiTool (Russo *et al*., 2018), which uses the WGCNA algorithm (Langfelder and Horvath, 2008) (A), monocle3 (Cao *et al*., 2019) (B) and fcoex (C). D) Comparison of the number of genes per module for each of the algorithms. E) Multi-hierarchical view of the *pbmc3k* dataset derived from module-based reclusterings. Colours indicate which of the 3 algorithms allowed inference of an upper class that grouped cells beyond the original Seurat

## Notes

### Competing Interest Statement

The authors have declared no competing interest.

